# Rheological transition driven by matrix makes cancer spheroids resilient under confinement

**DOI:** 10.1101/2022.02.09.479678

**Authors:** Tavishi Dutt, Jimpi Langthasa, U Monica, Satyarthi Mishra, Siddharth Bothra, Annapurna Vadaparty, Prosenjit Sen, Ramray Bhat

**Author notes:** Corresponding authorship.

## Abstract

Cancer metastasis through a confining peritoneal fluid microenvironment is mediated by spheroids: clusters of disseminated transformed cells. Ovarian cancer spheroids are frequently cavitated and their ‘blastuloid’ morphology is correlated with an extracellular matrix (ECM) coat. Here, we investigate the effects of such morphology on the mechanical integrity of confined cancer spheroids. Atomic force microscopy showed higher elastic modulus for blastuloid spheroids relative to their prefiguring non-lumen ‘moruloid’ counterparts. Subsequently, spheroids were flowed through microfluidic conditions mimicking peritoneal confinement. Traversing moruloids exhibited asymmetric cell flows during entry, often deformed and disintegrated through travel, and showed an incomplete- and slow shape recovery upon exit. In contrast, blastuloids traveled faster, exhibited rapid and efficient shape recovery upon exit, symmetric vector flows, and lesser disintegration. A multiscale computational model predicted higher intercellular adhesion and a dynamical lumen make blastuloids resilient. Although, E-cadherin overexpression in moruloids did not affect their resilience, blastuloid ECM-debridement decreased E-cadherin membrane localization, obliterated the lumen, and reversed the rheological properties of blastuloids to those typifying moruloids. The ECM-induced lumen therefore drives spheroidal transition from a labile viscoplastic to a resilient elastic state allowing them to survive spatially-constrained peritoneal flows.

## Introduction

The peritoneal cavity of a patient suffering from advanced epithelial ovarian cancer is filled with disseminated multicellular aggregates, commonly known as spheroids ^1^. These spheroids colonize abdominal organs leading to metastasis ^2^. Metastasizing spheroidal cancer cells frequently become resistant to chemotherapeutic drugs necessitating a rigorous investigation of mechanisms underlying their formation and stability. Spheroids obtained by tapping the malignant ascites of ovarian cancer patients show heterogeneous morphologies: some exhibit a dysmorphic ‘moruloid’ (mulberry-like) phenotype, and others show smooth compacted surfaces and an internal lumen, giving them a ‘blastuloid’ appearance ^3,4^. On the one hand, these diverse multicellular phenotypes could be aggregative consequences of phenotypically heterogeneous cell types that are shredded into the peritoneum. On the other hand, they could represent progressive stages of metastasis with moruloid phenotypes maturing into their blastuloid counterparts ^5^.

There is a burgeoning body of literature on biophysical investigations of tumorigenic cellular ensembles. Of these, most studies focus on the migrational dynamics of spheroidal or tumoroidal cells within stromal-like extracellular matrix (ECM) microenvironments ^6, 7, 8^. The dynamics of ECM confinement, local growth, and mesoscale processes such as adhesion and diffusive gradients of signaling and proteolysis are relevant to such contexts. However, in fluid microenvironments, the assembly of multicellular structures from suspended single cells likely employ distinct mechanisms ^9,10^. Although elegant theoretical models have been constructed recently to explain dynamical structural transitions, technical difficulties of efficiently imaging floating clusters have allowed few biophysical characterizations of spheroids ^11,3^. Notable experimental exceptions include efforts to mechanically analyze spheroids using micro-tweezers, wherein those constituted from breast cancer cells were found to be softer than from their untransformed controls ^12^, and investigations using cavitational rheology to determine the cortical tension in spheroids of HEK293 cells ^13^. A pertinent study by Panwhar and coworkers recently describes a high throughput approach using virtual liquid-bound channels to show that the stiffness of multicellular spheroids is an order of magnitude lower than that of cells that constitute them ^14^. Although these investigations have not studied temporal topological transitions between multicellular morphologies, they lay the foundation for such studies within fluid microenvironments.

Whereas detachment from adhesive substrata generate cellular stresses ^15^ , additional stresses are imposed on clusters due to movement through fluid spaces ^16, 17, 18^. Circulatory spaces such as the peritoneum have confining tunnels ^19^ and microcavities such as stomata ^20^ that cancer spheroids must negotiate safely for their survival and eventual metastasis.

In this paper, we have probed the mechanical properties of ovarian cancer spheroids upon their traversal through a spatially constraining microfluidic channel. Previous studies have validated microfluidic approaches to be particularly useful in analyzing the mechanical properties of biomaterials, such as localized forces like traction, and their recovery dynamics ^21,22^. We demonstrate that moruloid and blastuloid spheroids behave distinctly under constrictive flow. Our biophysical investigations in combination with multiscale modelling, reveal the importance of the spheroidal ECM in multicellular morphological transitions and provide insight into the sustained endurance of spheroids within spatially confined geometries of the peritoneal cavity.

## Results

### A cavitational blastuloid phenotype shows elastic behavior

Moruloid (lumen-less) and blastuloid (lumen-containing) spheroids form in a temporally successive progression from the ovarian cancer cell line OVCAR3 upon suspended cultivation in low adherent conditions (Fig. 1A; red and white represent fluorescent signals for F-actin and DNA respectively). We first probed for differences in their bulk mechanical behavior using atomic force microscopy (AFM). To stabilize spheroids for AFM, we suspended them on top of agar beds (schematic shown in Fig. 1Bi; see materials and methods for a detailed description of the agar preparation). We observed that blastuloids showed a higher mean elastic modulus (555 Pa) compared to moruloids (342 Pa) (Figure 1Bii). To further probe their mechanical properties under suspension, we subjected them to flow assays within a poly dimethyl siloxane (PDMS)-constructed microfluidic channel that would constrain their movement in terms of size (Fig. 1Ci). High speed videography showed that moruloids emerged from the constrictive channel by assuming a shape that was deformed relative to their entry shape (Fig. 1Cii; Video S1). We further quantified the deformation by measuring the aspect ratio of the leading edge of the spheroids as they exited the microfluidic channel. The aspect ratio was defined as

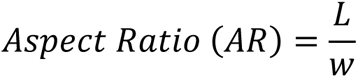

**Figure 1:**
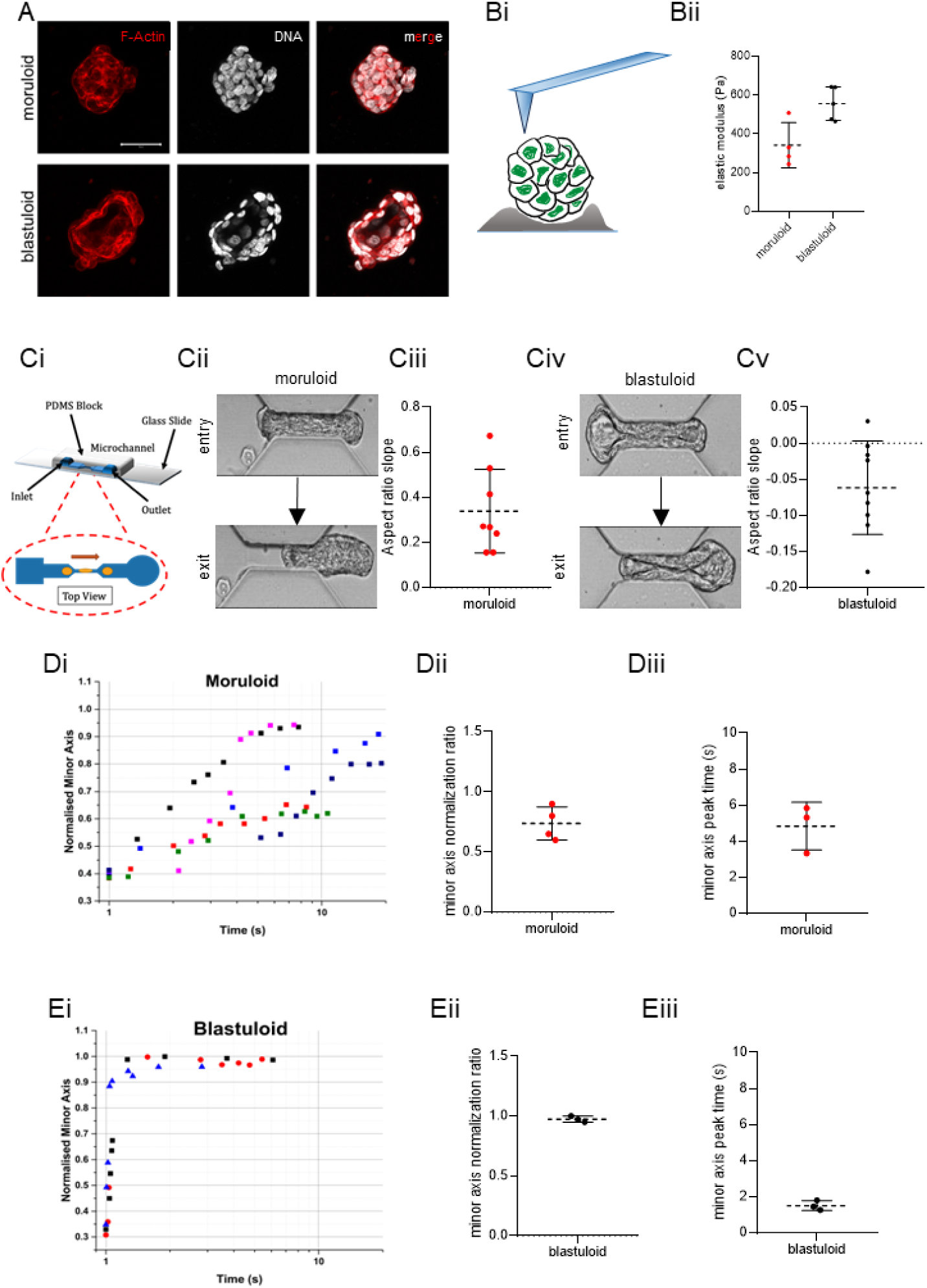
Ovarian cancer blastuloids relax efficiently and rapidly upon deformation. (A) Laser confocal micrographs of moruloid (top) and blastuloid (bottom) OVCAR3 spheroids showing maximum intensity projections of the fluorescence values representing F-actin (phalloidin; red) and DNA (DAPI; white). (Bi) Schematic depiction of atomic force microscopy performed on spheroids placed on agar beds. (Bii) Graph showing elastic moduli of moruloids (red dots) and blastuloids (black dots) (Ci) Schematic depiction of a microfluidic channel chip used for performing the flow experiments. (Cii) Snapshots of high-speed time lapse videography moruloids at the entry (top) and exit (bottom) of the channel (see also Video S1) (Ciii) Graph showing the aspect ratio slopes of exiting moruloids (see also Fig. S1) (Civ) Snapshots of high-speed time lapse videography blastuloids at the entry (top) and exit (bottom) of the channel (see also Video S2) (Cv) Graph showing the aspect ratio slopes of exiting blastuloids. (Di) Representative traces showing change in minor axis of exited relaxing moruloids (Dii) minor axis normalization ratio for exited relaxing moruloids (Diii) Time to peak for the minor axis of exited relaxing moruloids (Ei) Representative traces showing change in minor axis of exited relaxing blastuloids (Eii) minor axis normalization ratio for exited relaxing blastuloids (Eiii) Time to peak for the minor axis of exited relaxing blastuloids. Error bars represent mean+/- SD. Scale bars = 50 um. Refer to Supplementary Figures 1, 2 and 4 for more information.

where *w* is the minor axis and *L* is the major axis length of the protruding front-end of the spheroids (Fig. S1). These were quantified using a curve-fitting equation representing that of an ellipse (see Materials and Methods for further details). The parameters returned from the fit were used to obtain the aspect ratio for each individual protruded spheroid, for different time-points during its exit. Change in the aspect ratios was tracked as a function of time for representative clusters of each subset (Fig. S1).

The slopes were obtained by employing a simple linear fit on this curve. The mean slope for moruloids was found to be 0.33 (Fig. 1Ciii). In contrast, blastuloids were observed to regain their shape upon exiting the channel (Fig. 1Civ; Video S2). Consistent with these observations, the mean aspect ratio slope for blastuloids was -0.06, indicating the minor axis lengthened in proportion to the major axis as the spheroids progressively moved out of the channel (Fig. 1Cv). The difference in ingress and egress characteristics were also observed for non-cavitated (moruloid) and cavitated (blastuloid) spheroids derived from G1M2 ovarian cancer patient xenograft cells ((Fig. S2, see also Video S3 and S4) and from freshly harvested spheroids of a high grade serous ovarian cancer patient (Fig. S2, see also Video S5 and S6). We next traced the change in minor axis of spheroids normalized to their pre-entry values after spheroids had exited the channel to study the kinetics of their relaxation post deformation. Moruloids showed variable but incomplete extents of recovery (Fig. 1Di) with the mean normalized minor axis ratio at 0.8 (Fig. 1Dii) and its maxima achieved in an average time period of 4.8 sec (Fig. 1Diii). In contrast, blastuloids relaxed very quickly (Fig 1Ei) with the mean normalized minor axis ratio at 0.97 (Fig. 1Eii), plateauing within 1.5 sec (Fig. 1Eiii). We next asked whether the kinetics of entry and transit of such spheroids illuminate the differences in their mechanical behavior observed above.

### Constrictive traversal of blastuloid but not moruloid spheroids is size-autonomous

To verify how the motility of the spheroids through constrictive spaces would depend on their morphological parameters, we measured three time-metrics:

(i) **Entry Time**: defined as the time taken by the spheroid to fully enter the channel and calculated as the difference between the times when the leading edge first touches the channel, and the rear edge of the cluster first touches the channel (Fig. 2Ai).
(ii) **Travel Time:** defined as the time taken by the cluster to move through the channel, before the leading edge leaves the channel and calculated as the difference between transit time and entry time (Fig. 2 Aii).
(iii) **Transit time**: defined as the time a spheroid takes to pass through the channel and calculated as the difference between the times when the leading edge first touches the channel, and the rear edge of the cluster is just about to exit the channel (Fig. 2Aiii).

**Figure 2:**
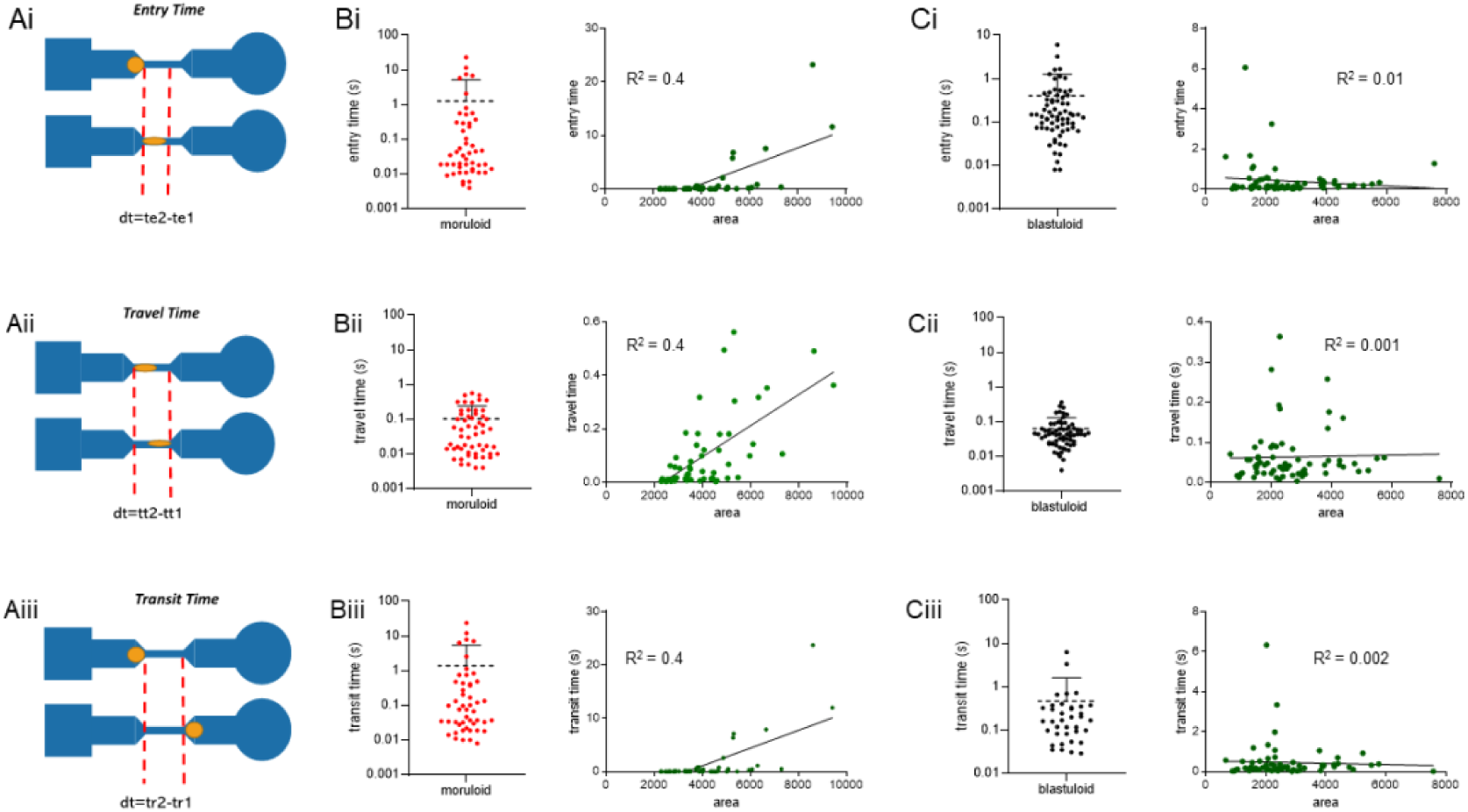
Ovarian cancer blastuloids travel through the channel faster than moruloid spheroids. (A) Schematic representation of three analyzed time metrics entry time (Ai), travel time (Aii), and transit time (Aiii). (B) Graphs showing entry time (Bi), travel times (Bii), and transit time (Biii) on the left and time-size correlation plots on the right for moruloids. (C) Scatter graphs showing entry time (Ci), travel times (Cii), and transit time (Ciii) on the left and time-size correlation plots on the right for blastuloids.

The mean entry time, travel time and transit time for moruloids was calculated to be 1.26 sec, 0.11 sec, and 1.37 sec, respectively (Fig. 2Bi-iii left). We also observed a moderate correlation of all temporal metrics with spheroidal size (R^2^ = 0.4) indicating bigger spheroids took longer to travel through the channel (Fig. 2Bi-iii right). The mean entry time, travel time and transit time for blastuloids were calculated to be 0.4 sec, 0.06 sec, and 0.46 sec (Fig. 2Ci-iii left). In addition, the entry, travel, and transit time for blastuloids were poorly correlated with spheroid size (R^2^ = 0.01, 0.001, and 0.002 respectively), indicating the mechanism by which blastuloids navigated the channel was relatively independent of their sizes (Fig. 2Ci-iii right). We next asked if the increased time for entry and travel observed for moruloids was due to rearrangements in intercellular contacts because of travel through constrictive spaces.

### Intercellular positional variation occurs within moruloid spheroids during entry into channel

To comprehend the flow of cells within spheroids when they traverse the channel, we used the PIVlab plugin in MATLAB to overlay morphologies with vectors (green) ^23,24^. We measured the normalized mean vector value: the ratio between the mean magnitudes of the vectors overlaid over the spheroidal interior in given areas, just above (A1), and below (A2), the portion that entered the tunnel (Fig. 3Ai). In the case of moruloid spheroids, we observed a standard deviation of 0.84 in the value (Fig. 3Aii), indicative of a heterogeneity in velocities of unjammed intraspheroidal cells as they moved into the channel (heterogeneity in vector magnitudes was also observed in ingressing G1M2 and ascites-harvested moruloids (Fig. S2)). As moruloids exited, the vectors overlaid on egressed cells were still aligned to each other; this is consistent with the deformation and retarded relaxation seen for moruloids in our earlier experiments (Fig. 3Aiii, see also Fig S2). The magnitudes of the normalized mean vector value for blastuloids showed a smaller standard deviation of 0.2 suggesting a synchronized advective cell movement without any intraspheroidal cellular unjamming (Fig. 3Bi-ii; similar behavior seen for G1M2 xenograft cell line and ascites -harvested spheroids in Fig S2). During exit, vectors overlayed on egressing cells diverged away from each other indicating the rapid relaxation and deformation recovery observed before (Fig.3Biii and Fig. S2).

**Figure 3:**
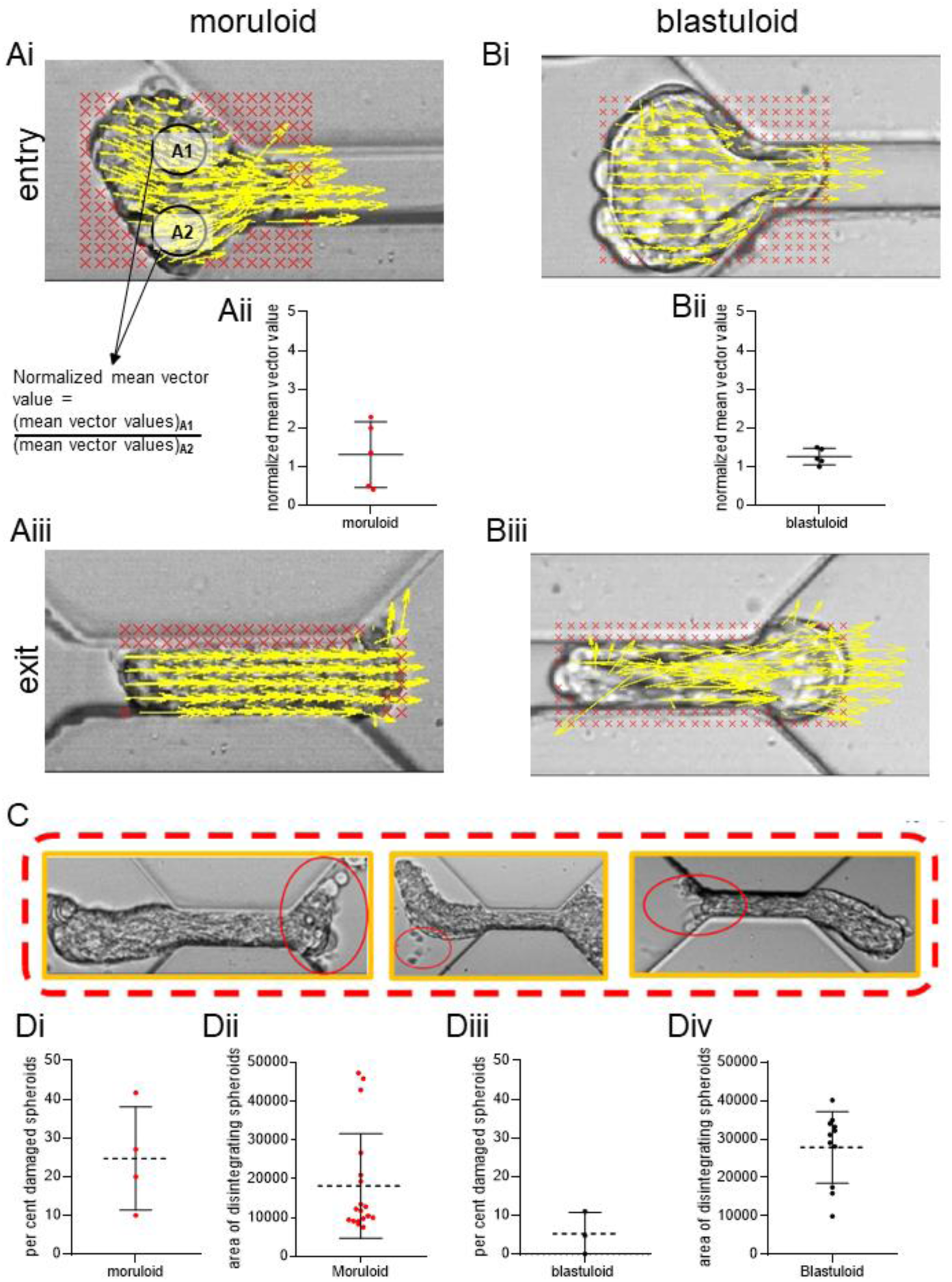
Ovarian cancer blastuloids exhibit minimal Intercellular rearrangement. (A) Particle image velocimetry (PIV) shows vectors sized based on magnitudes (yellow) overlaid on image of ingressing moruloids (Ai) with normalized mean vector values measured in areas A1 and A2 in the interior of spheroids (Aii). PIV vectors overlaid on image of egressing spheroids (Aiii). (B) Particle image velocimetry (PIV) shows vectors sized based on magnitudes (yellow) overlaid on image of ingressing moruloids (Bi) with normalized mean vector values measured in areas A1 and A2 in the interior of spheroids (Bii). PIV vectors overlaid on image of egressing spheroids (Biii). Red portions represent the masked areas not analyzed for flow. (C) Representative images showing cell detachment and disintegration in moruloid spheroids as they traverse the constrictive channel. (Di) Graph showing proportion of moruloids damaged due to traversal (Dii) Graph showing the area of moruloids that show damage upon constrictive traversal. (Diii) Graph showing proportion of blastuloids damaged due to traversal (Div) Graph showing the area of blastuloids that show damage upon constrictive traversal. Error bars represent mean+/- SD Scale bars = 50 um.

This led us to ask if the unjamming seen during entry of moruloid spheroids makes them vulnerable to cell detachment and spheroidal fractures. Upon close examination of the videomicrography of moruloid spheroids, we observed evidence of cell detachment and spheroidal damage and disintegration (Fig 3C). Extending our observations across a wide number of traversals, a mean proportion of 25% of moruloid spheroids were found to exhibit varying degrees of spheroidal damage (Fig. 3Di). Moreover, the mean size of moruloid spheroids beyond which disintegration was observed was 6534 sq µm (Fig. 3Dii). In contrast, disintegration or cell detachment was observed only in 5.5% of blastuloid spheroids (Fig. 3Diii). The mean size beyond which such damage was observed was 10019 sq µm (Fig. 3Div). These observations suggest that both spheroidal size and phenotype regulate morphological integrity in constrained movement. Our observations of intercellular detachment as an output of both these traits led us to investigate the effects of cell-cell adhesion on constrictive traversal of distinct morphologies.

### A multiscale model predicts lower cell adhesion in moruloid spheroids

Multiscale computational models have been used with a great deal of success to simulate experimentally elucidated observations and verify whether multicellular phenotypes can indeed emerge through the empirical interactions^25, 26,27^. Compucell3D represents one of the best-known simulation frameworks allowing an integration of Cellular Potts model-based solving of a Hamiltonian for contact energies between cellular constituents with PDE solvers of molecular constituents^28^. We therefore constructed multicellular models for the moruloid and blastuloid spheroids, which were driven through a constrictive channel in the simulation space, similar to our experimental setup. The input parameters that were tunable within such simulations were the contact energies (inverse of adhesion strength) between individual digital cells. We performed parameter scans for a range of intercellular contact energy values (refer to Table 2). We observed stable ‘digital moruloid’ formed for all the 4 (0-15) values of contact energies tested. For stable ‘digital blastuloid’ formation, the contact energy values between ‘core cells’ (which represent lumen), and that between the core cells and the peripheral cells (which represent the digital equivalents of biological cells) were fixed at 0 (high adhesion), while the contact energy values between peripheral cells were varied from 5 to 15 (see Table 2). For these stable spheroid conditions, we confirmed that upon exiting the simulated constrictive channel space ‘digital moruloid’ spheroids showed a sustained deformation and intercellular rearrangement (through observed intermixing of peripheral (blue) and central (green) cells of the spheroids) (Fig. 4A and Video S7). On the other hand, the digital blastuloid spheroids recovered their shape rapidly upon exit (Fig. 4B and Video S8). Insets of exiting digital moruloid and blastuloid spheroids showed intercellular rearrangements in the former but not the latter, showing consistency with our experimental findings (Fig. 4A and B insets to the right). Consistent with these observations, the aspect ratio of egressing digital moruloid morphologies when normalized to entry aspect ratio was found to be significantly higher than that for digital blastuloid counterparts (unpaired t-test p<0.0001; Fig. 4C). In fact, at lower values of intercellular adhesion (contact energy = 15), we also observed cell detachment and disintegration for both digital moruloid (Fig. 4D, Video S9) and blastuloid spheroids (Fig. 4E, Video S10). In agreement with the experimental observations, we found greater cell detachment in the case of digital moruloid spheroids (46.67%) than with the digital blastuloid spheroids (20%) (Fig. 4F). Upon increasing intercellular adhesion, a decrease in normalized aspect ratio was observed for digital moruloid spheroids, suggesting that cell-cell adhesion may regulate the unjamming-to-jamming transition of multicellular ensembles (one-way ANOVA p<0.0001; Fig. 4G).

**Table 1:** Table showing the parameters used for the AFM measurements.

|  |  |
| --- | --- |
| Nature of probe | Spherical |
| Diameter of probe | 5 $\mu\text{m}$ |
| Cantilever | Hydra 0.05nm |
| Force Constant | 45e-3 (N/m) |
| Sensitivity | 35 (V/ $\mu\text{m}$) |
| Forward Speed | 0.8 ($\mu\text{m}/\text{sec}$) |
| Backward Speed | 0.8 ($\mu\text{m}/\text{sec}$) |

**Table 2:** List of different parameter values used in the model. Multiple values assigned for some parameters correspond to the input variables and the values tested.

| Parameter | Value |
| --- | --- |
| Temperature ($T_m$) | 10 |
| **Contact energy Parameters** |  |
| Medium-Medium | 10 |
| Medium-Core | 10 |
| Medium-Peripheral | 10 |
| Medium-Chamber | 0 |
| Chamber-Chamber | 0 |
| Chamber-Core | 50 |
| Chamber-Peripheral | 50 |
| Core-Core | [0, 5, 10, 15] |
| Core-Peripheral | [0, 5, 10, 15] |
| Peripheral-Peripheral | [0, 5, 10, 15] |
| **Volume and surface area parameters for all cell types (unless specified otherwise)** |  |
| Target Volume | 25 |
| $\lambda_{vol}$ | 5 |
| Target surface area | 20 |
| $\lambda_{surf}$ | 5 |
| Target Volume (Core of blastuloid spheroid) | 400 |
| $\lambda_{vol}$ (Core of blastuloid spheroid) | 0.5 |
| Target surface area (Core of blastuloid spheroid) | 80 |
| $\lambda_{surf}$ (Core of blastuloid spheroid) | 0.5 |

**Figure 4:**
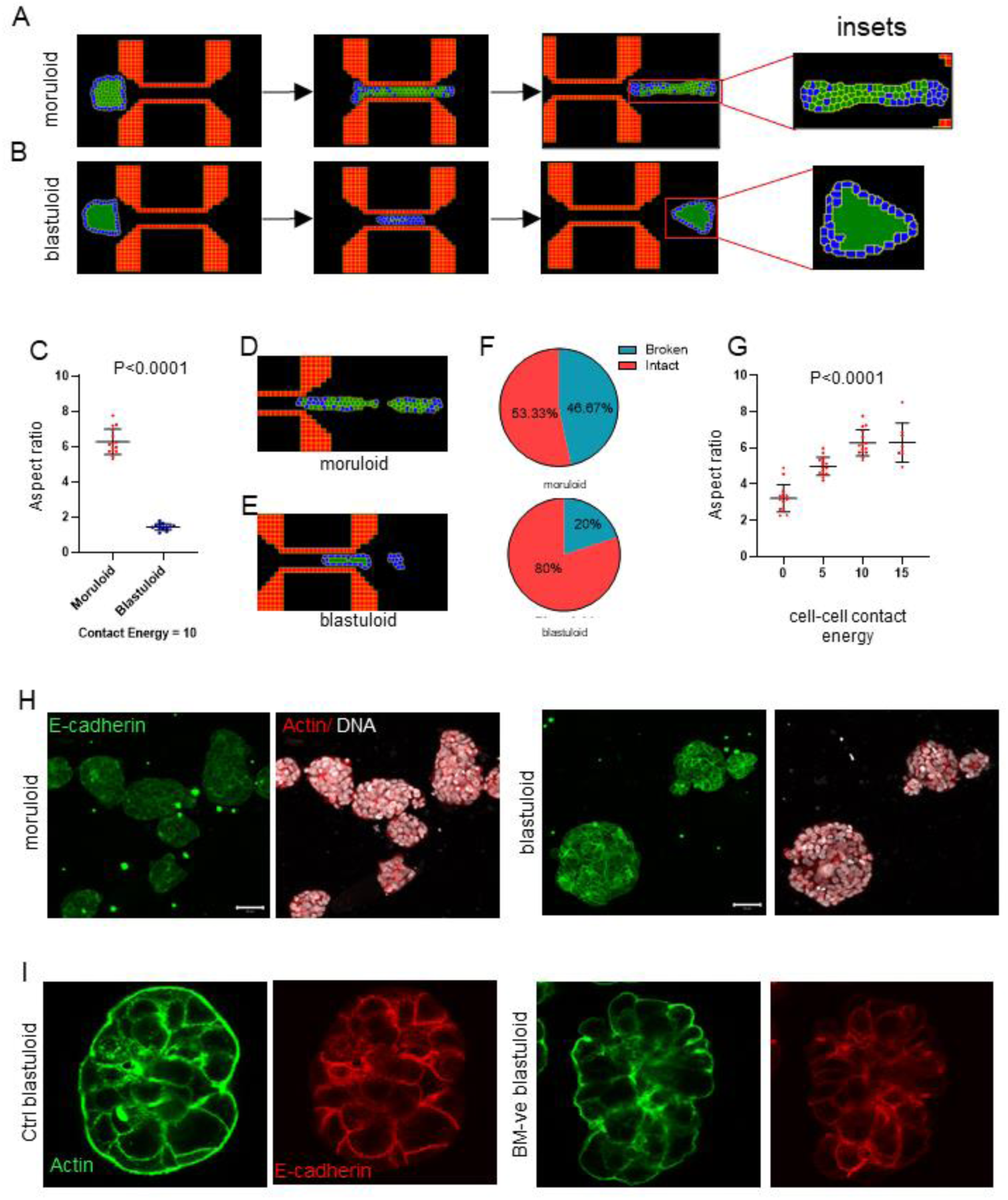
Ovarian cancer blastuloids show higher intercellular adhesion. (A, B) Snapshots of Compucell3D simulations of digital moruloid (A) and blastuloid (B) spheroids traversing through a spatially constrictive channel representing time points when the spheroids enter the channel (left) inside the channel (middle) and during exit from the channel (right). Insets show the intercellular arrangement within the digital spheroids upon exit. (C) Scatter graph showing the difference between aspect ratios of digital moruloid and blastuloid spheroids upon exit from the channel normalized to the same at entry. Unpaired Student’s t-test used to compute statistical significance. (D, E) Snapshot of a Compucell3D simulation of a digital moruloid (D) and blastuloid (E) spheroid showing disintegration and cell detachment upon traversal. (F) Pie-chart representing fraction of spheroid disintegration upon traversal for digital moruloid (top) and blastuloid (bottom) spheroids. (F) Scatter graph of the normalized aspect ratios of digital moruloid spheroids with different values of cell-cell contact energy used in the multiscale simulation. One-way ANOVA was used to compute statistical significance. (H) Laser confocal micrographs of moruloid and blastuloid OVCAR3 spheroids showing maximum intensity projection of the fluorescence values representing E-cadherin (green) and counterstaining for F-actin (phalloidin; red) and DNA (DAPI; white). (I) Laser confocal micrographs of blastuloid OVCAR3 spheroids (untreated control left) and upon Type IV collagenase treatment (right) showing middle stack of the fluorescence values representing E-cadherin (red) and counterstaining for F-actin (phalloidin; green). Error bars represent mean+/- SEM. Scale bars = 50 um and 20 um respectively.

Interepithelial adhesion is principally mediated through adherens junctions, established using homodimeric interactions between transmembrane cell adhesion molecules such as E-cadherin. The expression of E-cadherin has been shown to be under the regulation of Laminin 111, a principal constituent of epithelial basement membrane (BM) matrix. Our earlier demonstration of BM expression being characteristic of blastuloid ovarian cancer spheroids, our observation of a lesser propensity of blastuloids to show cell detachment and our theoretical prediction of an association of higher intercellular adhesion with lower aspect ratio led us to ask whether the blastuloid spheroids expressed comparatively greater levels of E-cadherin. Using fluorescent immunocytochemistry, we observed greater transmembrane signals for E-Cadherin in blastuloid spheroids compared with moruloids (Fig. 4H). Blastuloid architecture therefore differs from moruloid counterparts in terms of presence of the cavitation, an ECM coat and high intercellular E-cadherin localization. Upon ECM degradation, cavitation is lost. In addition, the intercellular staining of E-cadherin in blastuloids was found to be weaker than control untreated spheroids (Fig. 4I).

### Extracellular matrix regulates the mechanical behavior of blastuloid spheroids

To verify if an increased intercellular adhesion driven by higher E-cadherin localization contributed to the mechanical behavior of blastuloid spheroids, we overexpressed E-cadherin in OVCAR-3 cells (Fig S3) and subjected their moruloid spheroids to channel traversal. High speed videomicrography of E-cadherin overexpressing moruloid spheroids showed an unjammed cellular movement within spheroids during entry (Fig. 5Ai; Video S11). Like their plasmid control counterparts, E-cadherin overexpressing moruloid spheroids also exited in a deformed fashion suggesting slow relaxation kinetics (Fig. 5Aii).

**Figure 5:**
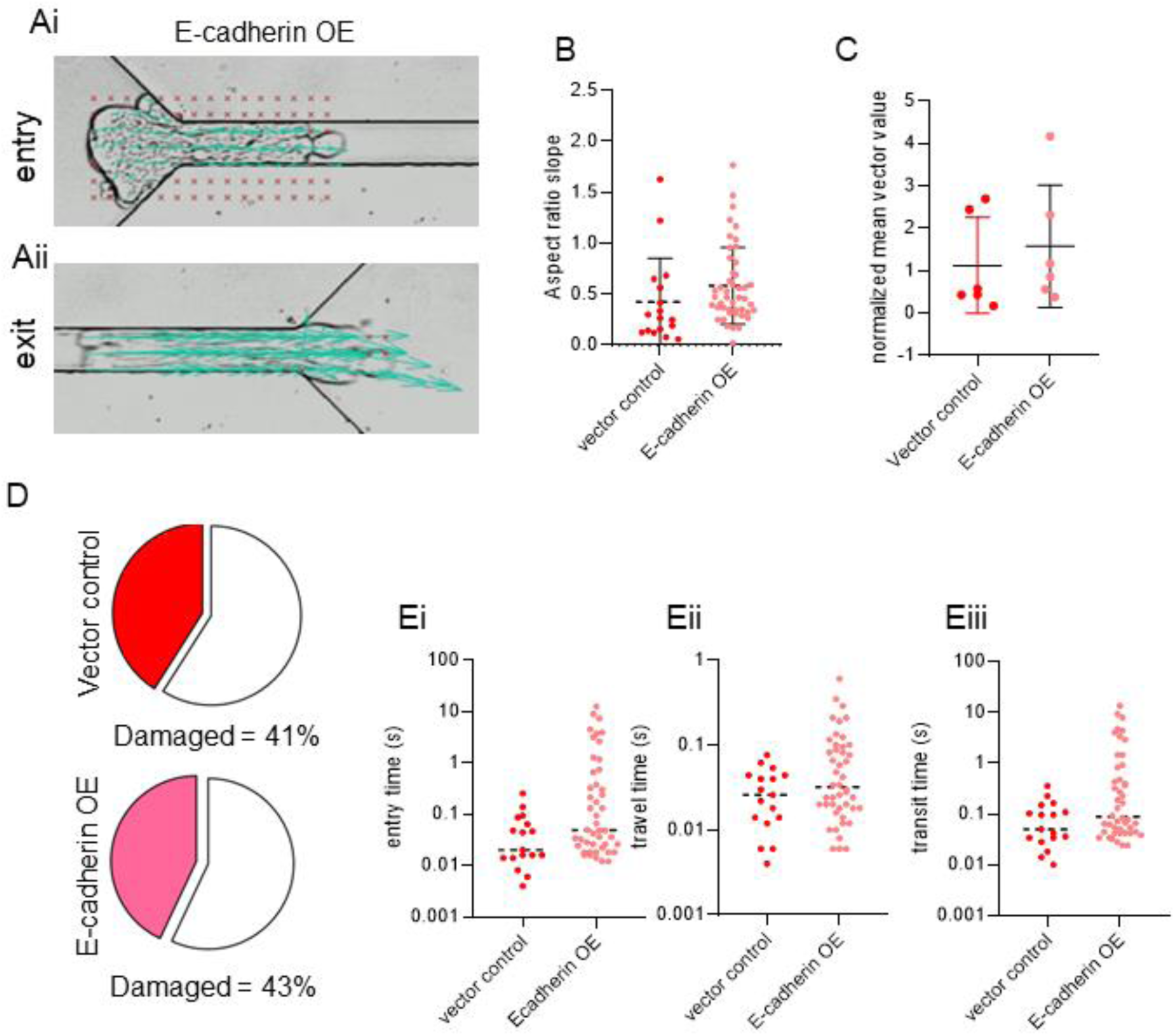
E-cadherin overexpression does not alter mechanical behavior of ovarian cancer moruloids. (A) Particle image velocimetry (PIV) shows vectors sized based on magnitudes (yellow) overlaid on images of E-cadherin-overexpressing moruloids at entry (Ai) and exit (Aii). Red portions represent the masked areas not analyzed for flow. (B) Graph showing the aspect ratio slopes of exiting E-cadherin overexpressing moruloids. Unpaired Student’s t-test used to compute statistical significance. (C) Graph showing normalized mean vector values measured in areas A1 and A2 in the interior of ingressing E-cadherin overexpressing moruloids shown in Ai (D) Pie charts showing proportion of damaged moruloids expressing empty vector (control) (red) and expressing E-cadherin (pink) (E) Graphs showing entry time (Ei), travel times (Eii), and transit time (Eiii) of vector and E-cadherin overexpressing moruloids. Unpaired Student’s t-test used to compute statistical significance. Error bars represent mean+/- SEM.

The aspect ratios of the spheroids at exit normalized to entry showed insignificant difference compared with plasmid controls (Fig. 5B unpaired t-test p = 0.1 File S1). The standard deviation of the normalized mean vector magnitudes for control and E-cadherin overexpressing moruloid spheroids entering the channel were 1.128 and 1.44, respectively (Fig. 5C). The proportion of spheroids which disintegrated upon exit were similar (41% and 43% damaged) for traversing control and E-cadherin overexpressing spheroids, respectively (Fig. 5D). In addition, we observed insignificant differences in the traversal temporal kinetics of E-cadherin-overexpressing spheroids compared with controls in terms of entry, travel, and transit times (Fig. 5Ei-iii unpaired t-test p = 0.08 for all three times File S1).

This led us to ask if ECM-driven morphogenesis (which included E-cadherin localization but also spheroidal cavitation) could drive the mechanical distinction between blastuloids and moruloids. Blastuloids were enzymatically debrided of their ECM coat using Type IV Collagenase (Fig. S4; green and white represent fluorescent signals for Type IV collagen and DNA, respectively). Atomic force microscopy showed that the mean elastic modulus of ECM-removed blastuloids was 337 Pa, which was significantly lower than blastuloids and closer to moruloid spheroids (one way ANOVA p = 0.008 (File S1); Fig 6A). A time lapse videographic examination showed that ECM-removed blastuloids showed deformation as they exited the microfluidic channel like moruloids (Fig. 6B) and their mean aspect ratio slope was 0.3 similar to moruloids and distinct from blastuloids (one way ANOVA p<0.0001 (File S1); Fig. 6C). We then assessed the relaxation kinetics of exited ECM-removed blastuloids and found them to recover only partially (Fig. 6Di-iii; maximum normalized minor axis = 0.8 (Fig. 6Dii) and median time for the minor axis to plateau was 7 sec) similar to moruloids (Fig. 6Diii) (one way ANOVA p = 0.01 and 0.02 respectively for the two metrics (File S1)). The temporal kinetics of ECM-removed blastuloids (median entry time = 2.8 sec (Fig 6Ei), mean travel time = 0.14 sec (Fig. 6Eii), and mean transit time = 2.9 sec (Fig. 6Eiii)) resembled that of moruloids and significantly greater than blastuloids (one way ANOVA p = 0.002, 0.01, and 0.002 respectively (File S1)). Except for travel time, both entry time and transit time showed moderate correlation with spheroidal size, resembling moruloid rather than blastuloid behavior (R^2^ = 0.22 for both and 0.001 for travel time; Fig. 6Fi-iii). Particle image velocimetry on ECM-removed blastuloids entering the channel revealed heterogeneous vector magnitudes across the entering spheroid portion (standard deviation of normalized mean vector values = 0.59 much higher than for blastuloids Fig. 6Gi-ii), smooth transit followed by an exit in which the vectors overlaid on exiting cells were aligned with each other indicative of a sustained deformation and delayed relaxation during egress (Fig. 6Gi). ECM removal in blastuloid spheroids also led to an increase in propensity for damage (42%) (Fig. 6Hi; one way ANOVA p = 0.03 File S1) with the maximum size at which disintegration observed being 1799 sq µm (Fig. 6Hii; one way ANOVA p < 0.001 File S1).

**Figure 6:**
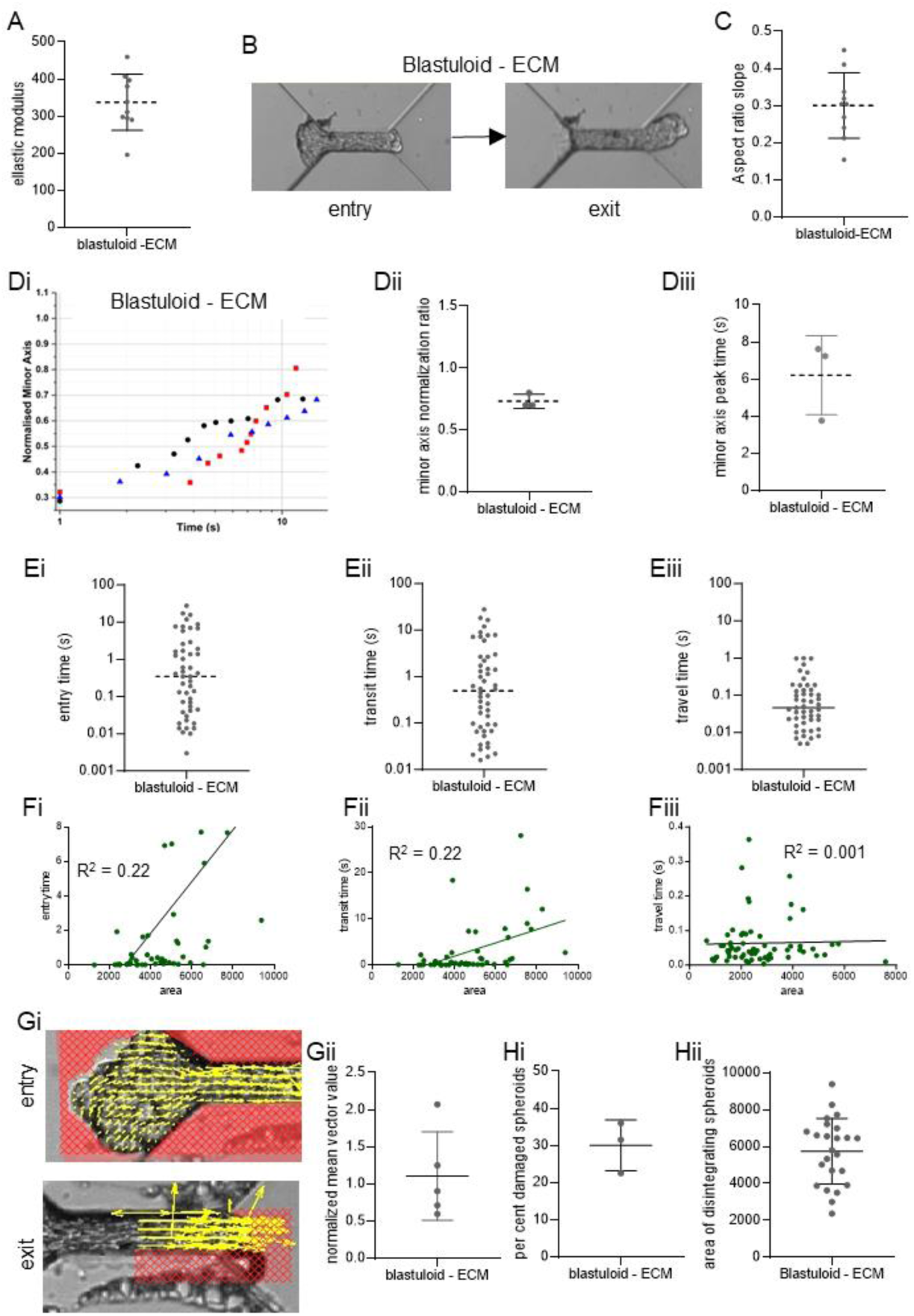
ECM removal in blastuloids results in mechanical behavior typical of moruloids. (A) Graph showing elastic moduli of blastuloids upon ECM removal. (B) Snapshots of high-speed time lapse videography blastuloids with ECM removal at the entry (left) and exit (right) of the channel (C) Graph showing the aspect ratio slopes of exiting blastuloids upon ECM removal (see also Fig S1) (Di) Representative traces showing change in minor axis of exited relaxing ECM-removed blastuloids (Dii) Minor axis normalization ratio for exited relaxing ECM-removed blastuloids (Diii) Time to peak for the minor axis of exited relaxing ECM-removed blastuloids. (E) Graphs showing entry time (Ei), travel times (Eii), and transit time (Eiii) of ECM-removed blastuloids. (F) Plots showing entry time-travel time- and transit time-correlations with spheroid sizes (Fi-iii) (Gi) Particle image velocimetry (PIV) shows vectors sized based on magnitudes (yellow) overlaid on images of ECM-removed blastuloids at entry (top) and exit (bottom). Red portions represent the masked areas not analyzed for flow. (Gii) Graph showing normalized mean vector values measured in ingressing ECM-removed blastuloids shown in Gi (Hi) (Diii) Graph showing proportion of ECM-removed blastuloids damaged due to traversal (Div) Graph showing the area of ECM-removed blastuloids that show damage upon constrictive traversal.

## Discussion

Our study is centered on an important observation: that the same cancer cell can form distinct multicellular architectures with unique mechanical properties. A spheroid that is constituted entirely of cells with no internal lumen exhibits deformability upon strain due to spatially constrictive travel. In addition, their shape recovery timescales are far slower than their deformation timescales indicating irreversible changes in their mechanical structure indicative of plastic behavior. Under strain they also show cellular disintegration. On the other hand, a lumen containing spheroid shows a jammed cellular organization with a low propensity for cellular detachment. Post deformation, their recovery timescales are much faster and complete, suggesting their morphology undergoes little change under constriction. Although this could be indicative of an elastic behavior of materials, their shorter entry times suggests that blastuloid spheroids are able to maintain their morphology by contextually reducing their luminal volume during entry. In fact, initial attempts to computationally model the blastuloid spheroids as impermeable elastic spheres resulted in longer entry times in simulations as well. This was mitigated upon simulating a dynamic deflation-inflation behavior of the digital blastuloid spheroids during traversal. The kinetics of lumen reinflation of these spheroids upon exiting the constriction suggests that the lumen acts as a dynamical volume whose magnitude is a function of the strain faced by the spheroid. Furthermore, the similarity between the behavior of blastuloid spheroids upon ECM removal and moruloid spheroids can be understood by how the basement membrane proteins mediate two crucial morphogenetic characteristics of blastuloid spheroids: maintenance of lumen and high intercellular adhesion. The lumen-less spheroids on the other hand are morphogenetically labile, owing to a loss of the external BM and lower intercellular adhesion. Therefore, the kinetics of regaining a more spheroidal shape occurs on a longer scale than for blastuloid spheroids. Further, the recovery is incomplete as is indicated by our PIV analysis, due to an irreversibly altered arrangement of the cells constituting them.

As moruloid spheroids differentiate into blastuloid spheroids, the BM induces a transition in cell migratory dynamics from an unjammed to a jammed state ^29,30^. It has not escaped our attention that another transition occurs concurrently: a mesenchymal to epithelial transition ^31,32^. The moruloid spheroids are associated with fibronectin expression, whereas we observe strong expression of E-cadherin in the blastuloid spheroids. Fibronectin and E-cadherin are classical markers of mesenchymal and epithelial states, respectively ^33^. Moreover, BM is also associated with the epithelial state of tissues ^34^. In addition, whereas a tendency to detach from a multicellular mass strongly characterizes a mesenchymal state, higher intercellular adhesion is typical of epithelial one ^35^. Taken together, the blastuloid-moruloid shift corresponds to a mesenchymal-to-epithelial transition (MET). Fredberg and colleagues have explored whether the transition between unjammed to jammed cell behavior and that between mesenchymal to epithelial states are conceptually non-congruent across biological examples ^35^. Our study seeks to bridge the two frameworks by showcasing how a limited set of proteins shift multicellular states both in terms of their polarity and motility.

Do spheroids encounter such spatially restricted spaces within the peritoneal cavity? Apart from the classical epiploic foramen that connects the lesser and greater sac, there are several peritoneal fossae as well as lymphatic stomata, wherein spheroids may potentially be subjected to constriction ^36^. Moreover, the peritoneal spaces are dynamic owing to constant motion within the surrounding organs, such as peristalsis. Such spatial restrictions could impinge on the rheology of diverse spheroidal morphologies and enhance the spectrum of their behaviors: cell detaching from the moruloid spheroids are better suited to micrometastatic colonization, whereas the jammed stable blastuloid spheroids could represent a niche suited for longer survival within the ascites. This is seen through the prolific accumulation of the cancer cells within the ascites of ovarian cancer patients.

Our study throws up engaging questions. On a mechanistic level, what is responsible for dynamical behavior of the blastuloid spheroids: the lumen, the BM or the higher intercellular adhesion? Given that all these processes are entangled phenomenologically, it is not easy to dissect the exact contribution of each of these to the ensemble properties. BM debridement and dysregulation of E-cadherin through genetic approaches would be employed in a temporally controlled manner in the future to dissect the origins of tissue elasticity. Multicellular ensembles have now been proposed to have greater potential for metastatic colonization than their unicellular counterparts. Could such potential be explained by their ability to withstand strain because of robustness to disintegration even in narrow peritoneal spaces? A broader generalization of our study to other cancers that employ both transperitoneal and vascular route of dissemination would shed light on the ubiquitous influence of mechanical regimes on disseminated cancer cell behavior.

## Materials and Methods

### Cell culture

Ovarian cancer cell lines used in this study are OVCAR3 (American Type Culture Collection; kind gift from Prof Rajan Dighe, Indian Institute of Science) and G1M2 (patient derived xenograft line; gift from Prof Sharmila Bapat, National Centre for Cell Sciences, India). These cell lines were maintained in Dulbecco’s Modified Eagle’s Medium - high glucose (AL007A; HiMedia) and Roswell Park Memorial Institute medium (AL162A; HiMedia) supplemented with 10-20 % FBS and recommended antibiotics in a humidified atmosphere of 95 % air and 5 % CO2 at 37°C. Cell identities were confirmed through STR analysis and were routinely tested for mycoplasma contamination.

### Spheroid culture and collagenase treatment

Spheroid were cultured in defined medium for 1 day to 7 days in suspension. Medium composition: DMEM: F12 (1:1) (HiMedia AT140) supplemented with 0.5 µg/ml hydrocortisone (Sigma-Aldrich, H0888), 250 ng/ml insulin (Sigma-Aldrich, I6634), 2.6 ng/ml sodium selenite (Sigma-Aldrich, S5261)), 27.3 pg/ml estradiol (Sigma-Aldrich, E2758), 5 µg/ml prolactin (Sigma-Aldrich L6520, 10 µg/ml transferrin (Sigma-Aldrich, T3309). 1.5 × 105 cells were seeded in polyHEMA coated 35 mm dishes for the experiment. Spheroids were collected from the cultures by centrifugation at 12000 rpm for 5 min. Collagenolysis of blastuloid spheroids was done by incubating with 600 U collagenase IV (17104-019 Gibco) for 24 hrs.

### Clinical samples

Ascites obtained from the peritoneal tap of patients with ovarian cancer was provided by Sri Sankara Cancer Hospital with due consent and ethical clearance from their Institutional Ethical Committee (SSCHRC/IEC4/015). Patient spheroids were cultured in tissue culture-treated polystyrene substrata/polyHEMA coated dish using DMEM (AL007A; HiMedia)—supplemented with 10–20% FBS (10270; Gibco) and antibiotics or with defined medium. Spheroids were then collected from the cultures by centrifugation at 0.1–0.2g for 5 min.

### Atomic Force Microscopy

A solution of noble agar dissolved in phosphate buffered saline (PBS) was heated in a microwave oven. The solution was immersed partially in a water bath and heated till the agar fully dissolved Different concentrations of this solution were tried resulting in optimal performance with 2% wt/vol. (0.04g of noble agar powder in 2mL of PBS).

The molten solution was poured on the edge of a pre-heated (to 37°C) 35mm petri-dish, and a freshly cleaned glass slide is used to smear the dispensed molten agar. Cells or spheroids were dispensed using a micro-pipette as big drops (∼100 μL) and allowed to rest for the agar to solidify and the cells to attach. 1mL of media or HEPES buffer is put in the final step and it is taken for the measurement. The AFM experiments were conducted using Park Systems XE-Bio AFM tool. The parameters used for the measurements are described in Table 2. The software used for analysis is the XEI software. The force-distance curves need to be converted to force-separation curves. However, this conversion is done by the XEI software which directly allows us to proceed from the force-separation curves.

The Young’s Modulus values are obtained from the software which uses a Hertzian Model to calculate the values, which is described by the following equation:

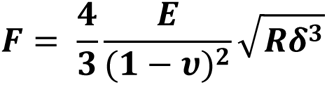

where,

is the applied force
is the Young’s Modulus
𝝊 is the Poisson’s Ratio
is the radius of the tip
δ is the indentation of the sample

## Gene expression analysis

Cells were lysed in the manufacturer’s recommended volume of Trizol (Takara, Japan). RNA was isolated by the chloroform-isopropanol extraction method. All reagents were molecular grade and purchased from Merck. Quantification of RNA yield was performed using the NanoDrop® ND-1000 UV-Vis Spectrophotometer (NanoDrop Technologies, USA). 1 μg of total RNA was reverse transcribed using Verso™ cDNA synthesis kit as per the manufacturer’s protocol (Thermo Scientific, AB-1453). Real-time PCR was performed with 1:2 diluted cDNA using SYBR green detection system (Thermo Fischer Scientific, F415L) and Rotorgene Q (Qiagen, 9001560). 18S rRNA 27 gene was used as the internal control for normalization. Relative gene expression was calculated using the comparative Ct method, and gene expression was normalized to unsorted cells. All the genes analyzed along with the sequence is mentioned in Appendix. Appropriate no template and no-RT controls were included in each experiment.

### Immunocytochemistry and Imaging

Spheroids were collected after centrifugation at 12000 rpm and fixed using 3.7% formaldehyde (24005; Thermo Fisher Scientific) at 4°C for 20 min. After one wash with PBS, the fixed spheroids were resuspended in PBS and put 10-20 µl of spheroids suspension to eight well chambered cover glass followed by placing on a dry bath at 37°C for 15–30 min for drying. Spheroids were then permeabilized using 0.5% Triton X-100 (MB031; HiMedia) for 2 h at RT. Using 3% BSA (MB083; HiMedia) prepared in 0.1% Triton X-100 solution blocking was achieved at RT incubated for 45 minutes. Primary antibody against E-cadherin (24E10 Cell Signaling Technology) was incubated overnight at 4°C which was followed by washes using 0.1% TritonX-100 in PBS (5 min × 3). Secondary antibody was incubated at RT for 2 h under dark conditions. DAPI (D1306; Thermo Fisher Scientific) was added to the samples and incubated for 15 minutes. Three washes were given after secondary antibody incubation step and after addition of DAPI. Images were captured in 20X and 40X using a Carl Zeiss LSM880 laser confocal microscope. Images were processed and analyzed using ZEN Lite software.

### Ellipse fitting algorithm for aspect ratio calculation

To quantify the visco-elastic or visco-plastic behavior of the spheroids, we opted to use Aspect Ratio of their ejection as a metric. It is obtained by fitting an ellipse to the protruding semi-circular shape coming out of the channel, for 4 different time points – front when the cluster is 25% exuding out of the channel to when it is almost 75%, on the verge of complete escape (supplementary). The ellipse fitting is done using a MATLAB code, to find out the major and minor axes of the hence obtained ellipse. Using those values, we calculated the aspect ratios of the protruding end. (Fig. S1). (https://www.mathworks.com/matlabcentral/fileexchange/15125-fitellipse-m).

Particle imaging velocimetry was performed using the app PIVLAB on MATLAB (Mathworks R2023a)^37^.

### Modeling framework

Compucell3D (CC3D) is a problem-solving environment based on the lattice-based GGH (Glazier-Graner-Hogeweg) model or CPM (Cellular Potts model) that was designed to model collective behavior of active matter^38^. This is done by calculating the Hamiltonian energy function at each simulation step. In the simulation lattice, each cell is represented by rectangular Euclidean lattice sites or pixels that share the same cell ID. The model evolves at each Monte Carlo Step (MCS), which consists of index-copy attempts of each pixel in the cell lattice. Calculation of the Hamiltonian (H) determines the allowed configuration and behavior of cells at each MCS.

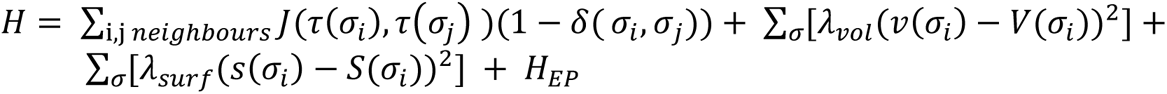

The Hamiltonian used in our model has four main contributors, which are affected by different properties of the cells. The first term in the energy function is the sum over all neighboring pairs of lattice sites *i* and *j* with associated contact energies (J) between the pair of cells indexed at those *i* and *j*. In this term, *i*, *j* denotes pixel index, σ denotes the cell index or ID, and τ denotes cell-type. The δ function ensures that only the σ_*i*_ ≠ σ_*j*_ terms are calculated (*i, j* belonging to the same cell will not be considered). Contact energies are symmetric in nature [𝐽(𝜏(𝜎_𝑖_), 𝜏(𝜎_𝑗_) ) = 𝐽(𝜏(𝜎_𝑗_), 𝜏(𝜎_𝑖_) )]. The contact energy between the two cells is considered inversely proportional to the adhesion between the two cells. The second term in the equation is a function of the volume constraint on the cell. λ_vol_(σ) denotes the inverse compressibility of the cell, *v*(σ) is the number of pixels in the cell (*volume*), and *V* (σ) is the cell’s target volume. The third term in the equation is a function of the surface area constraint on the cell, as the cells have fixed amounts of the cell membrane. For the cell σ, λ_surf_(σ) denotes the inverse membrane compressibility of the cell, *s*(σ) is the surface area of the cell, and *S*(σ) is the cell’s target surface area. The fourth term in the equation corresponds to the external potential applied to the center of mass of the cells to cause directional motion. 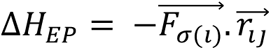 is the Hamiltonian for the external potential for a given MCS. For a cell σ, during an index copy attempt from *i* to *j,* the force vector is 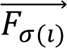 And the distance between the pixels *i* and *j* is 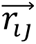 . The product of these two vectors along the right direction will lead to energy minimization and result in the movement of the cell along that direction.

The acceptation or rejection of the index copy attempt from pixel *i* to *j* depends on the change in the Hamiltonian (Δ*H*) due to the change in energy after the index copy attempt. When Δ*H* ≤ 0, the associated index-copy attempt will be successful, and the target pixels will be updated. So, the success probability is P = 1. When Δ*H* ≥ 0, the associated index-copy attempt will be successful following the Boltzmann probability function, with a probability of 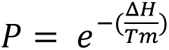 and it will be unsuccessful with a probability of *P′* = 1 – *P*. In the Boltzmann probability function, Δ*H* represents the calculated change in the overall Hamiltonian of the system between the system configuration at previous MCS and a specific system configuration at the current MCS. T_m_ relates to the effective membrane fluctuation for the cell and is kept at T_m_ = 10 in all the simulations.

A default dynamical algorithm known as modified Metropolis dynamics with Boltzmann acceptance function was used at each MCS to move the system towards a low-energy configuration as MCS increases. The term T_m_ can be considered temperature or magnitude of effective membrane fluctuations. Random movements of the pixels leading to different transition probabilities at each MCS mimics the stochasticity present in biological systems.

### Model components

We used a 450*200*1-pixel square lattice with a non-periodic boundary for all the simulations. Any model element required to participate through MCS pixel-copy attempts must be assigned a cell type. The model consisted of 4 different cell types: Chamber, Peripheral cells and Core cells, and the Medium. All cells were of dimension 5*5*1 in size, except for the core of the moruloid spheroids, of size 35*35*1. In an initial configuration, the chamber cells mimic the microfluidic chamber in the experimental setup, with the width to length aspect ratio maintained. These cells are “frozen” in the lattice, meaning the motility of the chamber cells is entirely restricted, and the movement of these cells to other lattice positions is not possible. The peripheral cells constituted the outer layer of the spheroids consisting of 32 cells. The core cells constituted the center of the spheroid. In the case of the moruloid spheroids, the properties of the core cells were similar to that of the peripheral cells. In the case of blastuloid spheroids, the core was made up of a single core cell. The lattices with no assigned cell type or, in other words, the free spaces were assigned as cell type ‘medium’ as a default by the CompuCell3D algorithm.

#### Contact energies (Differential Adhesion)

CompuCell3D requires setting interactions between all cell types in the model in terms of contact energies. Contact energy is inversely related to cell-cell adhesion. Higher contact energy between two cell types implies a higher contribution to the effective energy and lower probability of adhesion of the two cells. Since we have 4 different cell types, 10 different contact energy values must be assigned. These values were assigned based on previous literature ^38,39^. The contact energy values between core-core, peripheral-peripheral, and core-peripheral were considered input variables to observe the effect of morphology on the transition through the chamber (Table 2). In the case of moruloid spheroid, core-core, peripheral-peripheral, and core-peripheral values were the same as they all have similar properties.

#### External Potential

External potential was applied on the core and peripheral cells along the negative x-axis to enable movement of the spheroid through the chamber. After standardizing through parameter sensitisation, the value of the force vector was fixed at 6 for all simulations.

#### Compressibility

The compressibility of the core cells is different in the moruloid and the blastuloid spheroids to model the lumen in blastuloid spheroids. In the moruloid spheroids, the cells have higher inverse membrane compressibility fixed at 5, similar to the peripheral cells, making the cells stiffer. The core cell in the blastuloid spheroids, made to mimic the lumen, has lower inverse membrane compressibility fixed at 0.5 to make them more compressible and fluid-like.

#### Modelling deflation and inflation of blastuloid spheroids

The blastuloid spheroids were modelled to deflate when they met physical resistance while entering the narrow chamber. The velocity of the blastuloid spheroids reduced drastically when it met resistance while entering the chamber, and the core of the spheroids were modelled to shrink in size every time the velocity of the spheroid dropped considerably below the baseline velocity. The baseline velocity of these spheroids, averaged over a frequency of 100 MCS, was considered as the velocity at which they moved in the free space due to the constant force acting on them by means of the external potential. When the velocity of the core of the spheroids reduced to less than half its baseline velocity, the target volume and target surface area of the core of the spheroids was reduced, thereby mimicking deflation of the blastuloid spheroid. Similarly, on exiting the narrow chamber, the spheroid no longer experiences resistance and inflates. When the velocity of the core of the spheroids increased to twice its baseline velocity, the target volume and target surface area of the core was increased, to model the inflation of the spheroids. (https://github.com/MonicaUrd/Spheroid_tunnel_resistance_simulations.git).

### Microfluidic Chip Fabrication

A Piranha cleaned 4” Silicon wafer was spin coated with SU-2035 (Microchem) negative photoresist at 3000 rpm for 35 seconds to obtain a thickness of 75µm. Following a soft bake for – mins at - °C, it was UV-exposed using MJB-4 optical lithography tool through a 5” photomask containing the micropatterns. The wafer was then developed using SU-8 developer for 15 mins, followed by an IPA wash. It was coated with Teflon followed by a hard bake for 45 mins at 150°C.The pattern was transferred to a glass slide using soft lithography. PDMS was mixed with a curing agent in 10:1 ratio and poured on to the wafer with a mould. It was desiccated in vacuum for around 5 mins to remove any bubbles. After that it was heated for 45 mins on 110°C, and the hardened mould was peeled off once its cooled. Holes were punched for inlets and outlets using a biopsy punch, following which the PDMS block was plasma-bonded to an IPA cleaned glass slide after 1 min of treatment. The device was heated for 15 mins at 110°C to create a permanent bond.

### Experimental Setup

The sample was loaded into the device using a BD 1 ml syringe and microtubing. Using Chemyx Fusion high pressure pump, it was withdrawn (to avoid settling of clusters), at a rate of 1000 µl/hr. The flow was visualized and recorded using a high-speed camera at 1000 fps, with 10x (for the single channel) and 4X (for the 2-channel design) magnification lenses. Photron fast cam viewer (PFV-4) was used to view and analyze recorded videos. In the videos 1 µm corresponds to 1 pixel.

## Acknowledgments

We would like to acknowledge Shahid Hussain for help with spheroid preparation. This work was supported by the Wellcome Trust/DBT India Alliance Fellowship grant (IA/I/17/2/503312) awarded to RB. It was also supported by the John Templeton Foundation (#62220) and by the Department of Biotechnology, India [BT/909 PR26526/GET/119/92/2017 and BT/PR21962/NNT/28/1233/2017] to RB. PS acknowledges the Department of Biotechnology (DBT) and the Ministry of Electronics and Information Technology (MeitY). The opinions expressed in this paper are those of the authors and not those of the John Templeton Foundation. JL and TD acknowledge the Indian Institute of Science (IISc) for fellowship. MU and SB acknowledge KVPY for the student scholarship (X08010248, X113030157).

## Consent for publication

All the authors have provided their consent for publication of the manuscript.

## Availability of data and material

The raw data for all experiments and simulations will be made available upon request.

## Authors’ contributions

RB, PS and TD designed the experiments. TD, JL, SM, and SB performed the experiments. MU performed the simulations. RB, PS, TD, JL, MU, SM and SB analyzed the results of the experiments and simulations. AV performed the surgeries and provided the tissue samples. RB, PS, and TD wrote the manuscript. RB, PS, TD, JL, MU, SM, AV, and SB edited the manuscript.

## Supplementary Video Legends

Video S1: Transit of OVCAR3 moruloid through micro-channel.

Video S2: Transit of OVCAR3 blastuloid through micro-channel.

Video S3: Transit of G1M2 moruloid through micro-channel.

Video S4: Transit of G1M2 blastuloid through micro-channel.

Video S5: Transit of patient moruloid through micro-channel.

Video S6: Transit of patient blastuloid through micro-channel.

Video S7: Simulation of transit of a digital moruloid through a micro-channel

Video S8: Simulation of transit of a digital blastuloid through a micro-channel

Video S9: Simulation of transit of a digital moruloid with lower cell-cell adhesion through a micro-channel

Video S10: Simulation of transit of a digital blastuloid with lower cell-cell adhesion through a micro-channel

Video S11: Transit of OVCAR3 ECM-removed blastuloid through micro-channel.

## References

1. Lengyel E. Ovarian cancer development and metastasis. The American journal of pathology. 2010;177(3):1053–1064. doi:10.2353/ajpath.2010.100105

2. Shield K, Ackland ML, Ahmed N, Rice GE. Multicellular spheroids in ovarian cancer metastases: Biology and pathology. Gynecologic oncology. 2009;113(1):143–148. doi:10.1016/j.ygyno.2008.11.032

3. Langthasa J, Sarkar P, Narayanan S, Bhagat R, Vadaparty A, Bhat R. Extracellular matrix mediates moruloid-blastuloid morphodynamics in malignant ovarian spheroids. Life Science Alliance. 2021;4(10). doi:10.26508/lsa.202000942

4. Koike C, McKee TD, Pluen A, et al. Solid stress facilitates spheroid formation: potential involvement of hyaluronan. British journal of cancer. 2002;86(6):947—953. doi:10.1038/sj.bjc.6600158

5. Goyeneche A, Lisio MA, Fu L, et al. The Capacity of High-Grade Serous Ovarian Cancer Cells to Form Multicellular Structures Spontaneously along Disease Progression Correlates with Their Orthotopic Tumorigenicity in Immunosuppressed Mice. Cancers. 2020;12(3). doi:10.3390/cancers12030699

6. Kopanska KS, Alcheikh Y, Staneva R, Vignjevic D, Betz T. Tensile Forces Originating from Cancer Spheroids Facilitate Tumor Invasion. PloS one. 2016;11(6):e0156442. doi:10.1371/journal.pone.0156442

7. Pally D, Pramanik D, Hussain S, et al. Heterogeneity in 2,6-Linked Sialic Acids Potentiates Invasion of Breast Cancer Epithelia. ACS central science. 2021;7(1):110–125. doi:10.1021/acscentsci.0c00601

8. McKenzie AJ, Hicks SR, Svec K V, Naughton H, Edmunds ZL, Howe AK. The mechanical microenvironment regulates ovarian cancer cell morphology, migration, and spheroid disaggregation. Scientific reports. 2018;8(1):7228. doi:10.1038/s41598-018-25589-0

9. Duclut C, Sarkar N, Prost J, Jülicher F. Fluid pumping and active flexoelectricity can promote lumen nucleation in cell assemblies. Proceedings of the National Academy of Sciences. 2019;116(39):19264–19273. doi:10.1073/pnas.1908481116

10. Burleson KM, Casey RC, Skubitz KM, Pambuccian SE, Oegema TRJ, Skubitz APN. Ovarian carcinoma ascites spheroids adhere to extracellular matrix components and mesothelial cell monolayers. Gynecologic oncology. 2004;93(1):170–181. doi:10.1016/j.ygyno.2003.12.034

11. Duclut C, Prost J, Jülicher F. Hydraulic and electric control of cell spheroids. Proceedings of the National Academy of Sciences. 2021;118(19). doi:10.1073/pnas.2021972118

12. Jaiswal D, Cowley N, Bian Z, Zheng G, Claffey KP, Hoshino K. Stiffness analysis of 3D spheroids using microtweezers. PloS one. 2017;12(11):e0188346. doi:10.1371/journal.pone.0188346

13. Blumlein A, Williams N, McManus JJ. The mechanical properties of individual cell spheroids. Scientific reports. 2017;7(1):7346. doi:10.1038/s41598-017-07813-5

14. Panhwar MH, Czerwinski F, Dabbiru VAS, et al. High-throughput cell and spheroid mechanics in virtual fluidic channels. Nature Communications. 2020;11(1):2190. doi:10.1038/s41467-020-15813-9

15. Gilmore AP. Anoikis. Cell Death & Differentiation. 2005;12(S2):1473–1477. doi:10.1038/sj.cdd.4401723

16. Hyler AR, Baudoin NC, Brown MS, et al. Fluid shear stress impacts ovarian cancer cell viability, subcellular organization, and promotes genomic instability. PLOS ONE. 2018;13(3):e0194170-.

17. Zhao R, Afthinos A, Zhu T, et al. Cell sensing and decision-making in confinement: The role of TRPM7 in a tug of war between hydraulic pressure and cross-sectional area. Science Advances. 2019;5(7). doi:10.1126/sciadv.aaw7243

18. Follain G, Herrmann D, Harlepp S, et al. Fluids and their mechanics in tumour transit: shaping metastasis. Nature Reviews Cancer. 2020;20(2):107–124. doi:10.1038/s41568-019-0221-x

19. Bushati M, Rovers KP, Sommariva A, et al. The current practice of cytoreductive surgery and HIPEC for colorectal peritoneal metastases: Results of a worldwide web-based survey of the Peritoneal Surface Oncology Group International (PSOGI). European Journal of Surgical Oncology. 2018;44(12):1942–1948. doi:10.1016/j.ejso.2018.07.003

20. Wassilev W, Wedel T, Michailova K, Kühnel W. A scanning electron microscopy study of peritoneal stomata in different peritoneal regions. Annals of anatomy = Anatomischer Anzeiger : official organ of the Anatomische Gesellschaft. 1998;180(2):137–143. doi:10.1016/S0940-9602(98)80013-7

21. Nyberg KD, Hu KH, Kleinman SH, Khismatullin DB, Butte MJ, Rowat AC. Quantitative Deformability Cytometry: Rapid, Calibrated Measurements of Cell Mechanical Properties. Biophysical Journal. 2017;113(7):1574–1584. 10.1016/j.bpj.2017.06.073

22. Breslauer DN, Lee PJ, Lee LP. Microfluidics-based systems biology. Molecular bioSystems. 2006;2(2):97-112. doi:10.1039/b515632g

23. Khopkar AR, Aubin J, Xuereb C, Le Sauze N, Bertrand J, Ranade V V. Gas−Liquid Flow Generated by a Pitched-Blade Turbine: Particle Image Velocimetry Measurements and Computational Fluid Dynamics Simulations. Industrial & Engineering Chemistry Research. 2003;42(21):5318–5332. doi:10.1021/ie020954t

24. Cai S, Liang J, Gao Q, Xu C, Wei R. Particle Image Velocimetry Based on a Deep Learning Motion Estimator. IEEE Transactions on Instrumentation and Measurement. 2020;69(6):3538–3554. doi:10.1109/TIM.2019.2932649

25. Popławski NJ, Shirinifard A, Swat M, Glazier JA. Simulation of single-species bacterial-biofilm growth using the Glazier-Graner-Hogeweg model and the CompuCell3D modeling environment. Mathematical biosciences and engineering : MBE. 2008;5(2):355–388. doi:10.3934/mbe.2008.5.355

26. Ilina O, Gritsenko PG, Syga S, et al. Cell-cell adhesion and 3D matrix confinement determine jamming transitions in breast cancer invasion. Nature cell biology. 2020;22(9):1103–1115. doi:10.1038/s41556-020-0552-6

27. Ramos CH, Rodríguez-Sánchez E, Angel JAAD, et al. The environment topography alters the way to multicellularity in Myxococcus xanthus. Science advances. 2021;7(35). doi:10.1126/sciadv.abh2278

28. Pramanik D, Jolly MK, Bhat R. Matrix adhesion and remodeling diversifies modes of cancer invasion across spatial scales. Journal of theoretical biology. 2021;524:110733. doi:10.1016/j.jtbi.2021.110733

29. Kang W, Ferruzzi J, Spatarelu CP, et al. A novel jamming phase diagram links tumor invasion to non-equilibrium phase separation. iScience. 2021;24(11):103252. doi:10.1016/j.isci.2021.103252

30. Saitoh K, Hatano T, Ikeda A, Tighe BP. Stress Relaxation above and below the Jamming Transition. Physical review letters. 2020;124(11):118001. doi:10.1103/physrevlett.124.118001

31. Theveneau E, Mayor R. Neural crest migration: interplay between chemorepellents, chemoattractants, contact inhibition, epithelial-mesenchymal transition, and collective cell migration. Wiley interdisciplinary reviews Developmental biology. 2012;1(3):435–445. doi:10.1002/wdev.28

32. Bernardi Y, Strobl-Mazzulla PH. What we can learn from embryos to understand the mesenchymal-to-epithelial transition in tumor progression. The Biochemical journal. 2021;478(9):1809—1825. doi:10.1042/bcj20210083

33. Nowak E, Bednarek I. Aspects of the Epigenetic Regulation of EMT Related to Cancer Metastasis. Cells. 2021;10(12). doi:10.3390/cells10123435

34. Pozzi A, Yurchenco PD, Iozzo R V. The nature and biology of basement membranes. Matrix biology : journal of the International Society for Matrix Biology. 2017;57–58:1-11. doi:10.1016/j.matbio.2016.12.009

35. Le Bras GF, Taubenslag KJ, Andl CD. The regulation of cell-cell adhesion during epithelial-mesenchymal transition, motility and tumor progression. Cell adhesion &amp; migration. 2012;6(4):365—373. doi:10.4161/cam.21326

36. Thomas JM, Van Fossen K. Anatomy, Abdomen and Pelvis, Foramen of Winslow (Omental, Epiploic). StatPearls Publishing, Treasure Island (FL); 2021.

37. Thielicke W, Sonntag R. Particle Image Velocimetry for MATLAB: Accuracy and enhanced algorithms in PIVlab. Journal of Open Research Software. 2021;9(1):12. doi:10.5334/jors.334

38. Swat MH, Thomas GL, Belmonte JM, Shirinifard A, Hmeljak D, Glazier JA. Multi-scale modeling of tissues using CompuCell3D. Methods in cell biology. 2012;110:325–366. doi:10.1016/B978-0-12-388403-9.00013-8

39. Steinkamp MP, Winner KK, Davies S, et al. Ovarian tumor attachment, invasion, and vascularization reflect unique microenvironments in the peritoneum: insights from xenograft and mathematical models. Frontiers in oncology. 2013;3:97. doi:10.3389/fonc.2013.00097

