## Supplementary Figures for "Rheological transition driven by matrix makes cancer spheroids resilient under confinement"

S1

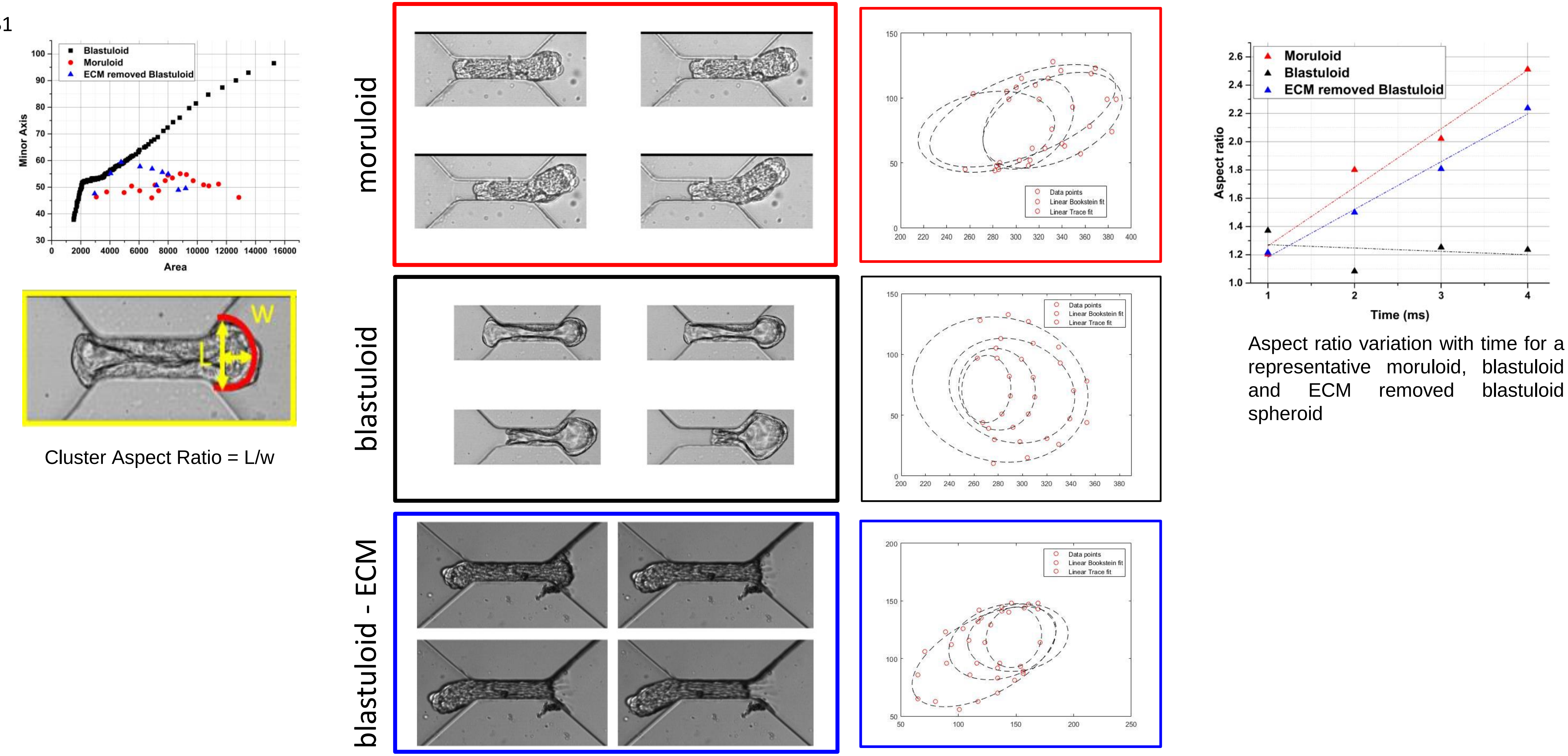


Cluster Aspect Ratio = L/w

Aspect ratio variation with time for a representative moruloid, blastuloid and ECM removed blastuloid spheroid

Supplementary figure 1: (Left top) Scatter plot showing minor axis evolution with area of the exiting cluster for a representative moruloid (red), blastuloid (black) and ECM-removed blastuloid. (Left bottom) Representative image showing Cluster Aspect Ratio calculated as L/w. Red curve represents the protruding portion used for ellipse fitting. (Middle) Calculation of aspect ratio for the high-speed images, taken at 4 timepoints during the exit of a representative moruloid, blastuloid and ECM-removed blastuloid spheroid; The ellipse is fit to the protruding edge using MATLAB and the obtained curves for all the 4 images (for each population) are shown next to the snapshots as dotted ellipses. (C) Aspect ratio variation with time for a representative moruloid, blastuloid and ECM-removed blastuloid spheroid. The 4 datapoints are fit using a linear fit and subsequent equation gives the slope

S2

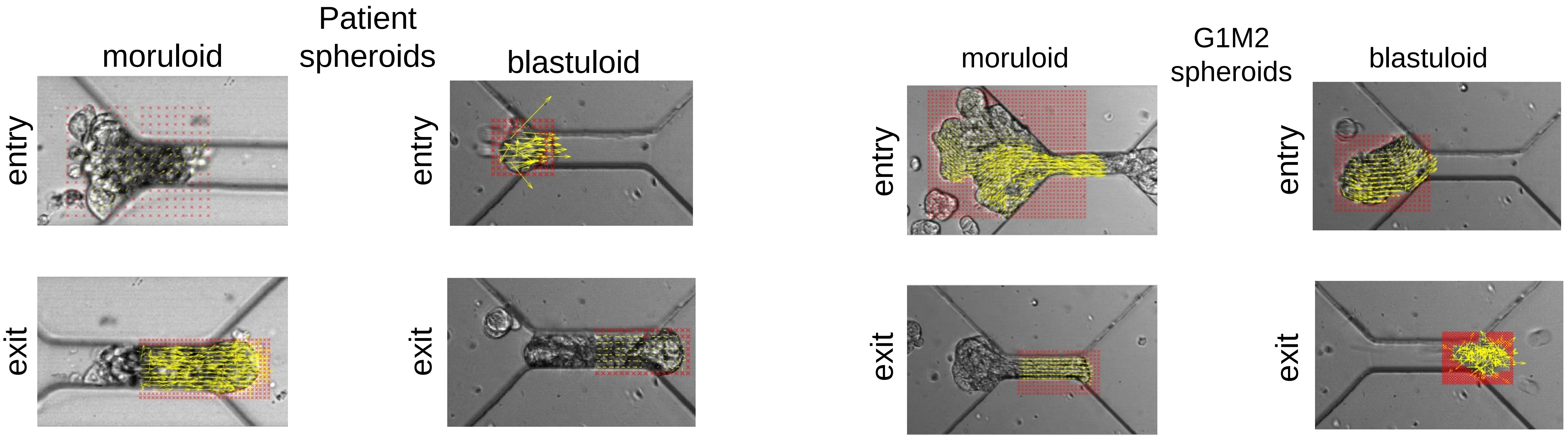


Supplementary Figure 2: Particle image velocimetry (PIV) shows vectors sized based on magnitudes (yellow) overlaid on image of ingressing moruloids (first from left) and blastuloids (second from left) from a patient with high grade serous ovarian cancer with ascites and from moruloids (second from right) and blastuloids (right most) from the G1M2 patient xenograft cell line. Top row shows images for spheroids during entry and the bottom row shows the images for spheroids during exit. Red portions represent the masked areas not analyzed for flow.

S3

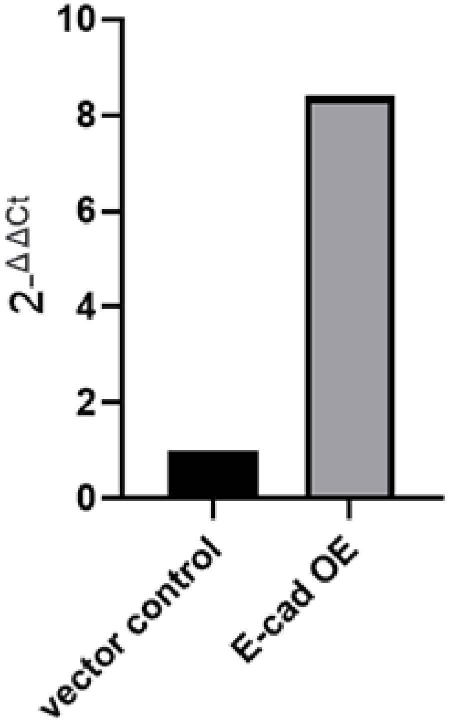


Supplementary Figure 3: qRT-PCR showing relative expression of E-Cadherin mRNA in cells which were transduced lentivirally with empty plasmid (vector control left) and human E-Cadherin whole length gene (right).

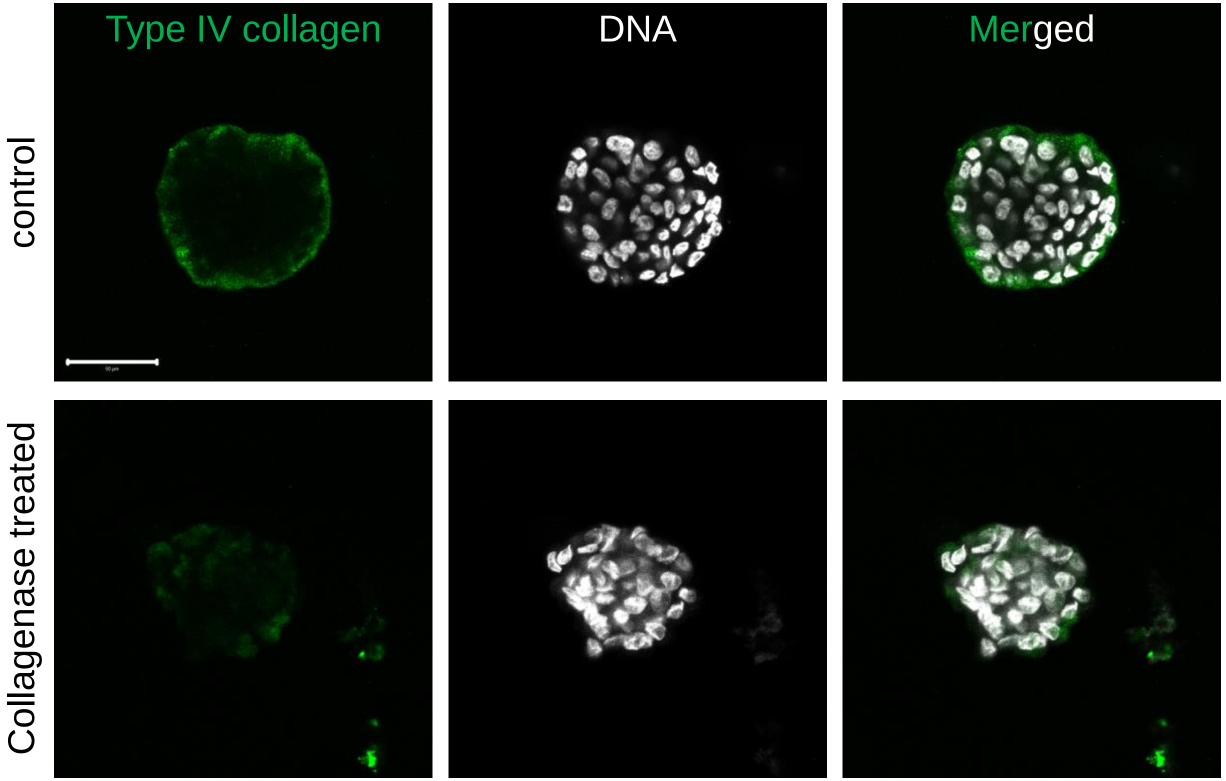


Supplementary Figure 4: Laser confocal micrographs of blastuloid OVCAR3 spheroids (top) and ECM-removed blastuloid OVCAR3 spheroids (bottom) showing mid stack of the fluorescence values representing Type IV collagen (green) and counterstaining for DNA (DAPI; white). Scale bars = 50 μm.
